# Diabetes-associated *MYT1* and *ST18* regulate human beta-cell insulin secretion and survival via other diabetes-risk genes

**DOI:** 10.1101/2025.02.24.639737

**Authors:** Ruiying Hu, Nala Hamilton, Yu Wang, Xin Tong, Mahircan Yagan, Prasanna K. Dadi, Cristina Harmelink, Teri D. Doss, Jinhua Liu, Yanwen Xu, Alan J. Simmons, Ken S. Lau, Roland Stein, Appakalai N. Balamurugan, Irina Kaverina, Kathryn C. Coate, David A. Jacobson, Qi Liu, Guoqiang Gu

## Abstract

**Aims/hypothesis:** Genetic and environmental factors work together to cause islet beta-cell failure, leading to type 2 diabetes (T2D). How these factors are integrated to regulate beta cells remains largely unclear. Based on our previous findings that the family of Myelin transcription factors (MYT1, MYT1L, and ST18) prevents mouse beta-cell failure by repressing the overactivation of stress response, their regulation by obesity-related nutrition signals in human beta cells, and their association with T2D, we postulate that these factors prevent human beta-cell failure under normal physiology and obesity-related stress.

**Methods:** *MYT1* or *ST18* were knocked down in primary human beta cells using shRNA. Beta-cell survival, secretory function, and gene expression were examined after islet cells were cultured *in vitro* or xenotransplanted into mice under normal or obesity-related stress.

**Results:** In culture, *MYT1-*knockdown (KD) caused beta-cell death, while *ST18-*KD compromised glucose-stimulated insulin secretion. Under obesity-induced stress as xenotransplants, *ST18-*KD also caused beta-cell death. Accordingly, *MYT1-*KD deregulated several genes and genesets in cell death and cellular stress response, while *ST18*-KD deregulated those regulating stress response, mitochondria, and ion channels. Corresponding to these gene expression changes, *ST18*-KD reduces glucose-stimulated Ca^2+^ influx in beta cells. In addition, the *MYT1-* and *ST18*-regulated genes are enriched for T2D-associated loci, with an enrichment of 2.05-fold relative to random distribution.

**Conclusions/interpretation:** The MYT TFs complement each other to integrate genetic and environmental factors to prevent beta-cell failure and T2D, with their major effects exerted on beta-cell viability and/or Ca^2+^ influx.

**Graphical Abstract:** Nutrient-responsive transcription factors MYT1 and ST18 regulate human beta-cell survival and secretory functions. Under low metabolic stress, MYT1 regulates cell survival while ST18 regulates insulin secretion. Under high metabolic stress, ST18 also regulates beta-cell survival. MYT1, at least partly, regulates cell survival through stress-related apoptotic processes, while ST18 regulates insulin secretion via Ca^2+^ influx.

**Research in context summary:** *Background:*

- The myelin transcription factors (MYT TFs, including MYT1, MYT1l, and ST18) prevent mouse beta-cell failure by depressing the overactivation of stress-response genes.
- SNPs in all three *MYT* loci are associated with human type 2 diabetes.
- The expression and nuclear localization of MYT1 and ST18 were increased in primary human beta cells under acute metabolic stress but downregulated in type 2 diabetes.
- The co-knockdown of *MYT1*, *MYT1L*, and *ST18* in a human beta-cell line resulted in apoptosis.

*Key question:*

- How does MYT1 or ST18 regulate human beta-cell function and survival?

*New findings:*

- MYT1 prevents human beta-cell death under normal and metabolic stress conditions, corresponding to deregulation of a few genes involved in cell death under cellular stress.
- ST18 promotes insulin secretion under normal physiological conditions by regulating Ca^2+^ influx and prevents beta-cell death under metabolic stress.
- The MYT1- and ST18-regulated genes are enriched for type 2 diabetes-risk loci.

*Clinical impact:*

- Regulators of the MYT-TF activities could be explored to delay/prevent beta-cell failure and the development of type 2 diabetes.

## Introduction

Type 2 diabetes (T2D) develops when beta cells cannot secrete sufficient insulin to maintain whole-body glucose homeostasis. Both genetic predisposition and non-genetic influences contribute to T2D. Hundreds of genetic loci have been associated with T2D risk (1), most of which regulate beta-cell dysfunction, loss of identity, or death (i.e., beta-cell failure) (2,3). Environmental factors exacerbate the problem by overwhelming beta cells with excessive workload or by causing the accumulation of stressor molecules (4). How beta cells integrate these factors to prevent this failure remains largely unknown.

A key risk factor for beta-cell failure is obesity, which induces insulin resistance (5–7). To compensate, beta cells increase insulin output in response to the high glucose and free fatty acid levels that accompany obesity. However, sustained high insulin output can exhaust the beta cells (6,8). High levels of glucose metabolism, required for glucose-stimulated insulin secretion (GSIS), promote insulin biosynthesis. This co-produces unfolded proinsulin in the ER that, if not cleared, induces ER stress and dysfunction (9). High glucose and free fatty acid metabolism in beta cells also produce reactive oxygen species and/or toxic lipid metabolites (10). These products, at high levels, cause oxidative stress and mitochondrial dysfunction while also exacerbating ER stress. Henceforth, beta-cells activate the stress response to maintain homeostasis (11,12). However, stress responses can persist too long or be overly activated, reducing essential beta-cell proteins or inducing proapoptotic genes (5,13). Thus, precisely regulating stress response is critical to prevent beta-cell failure (14).

The myelin transcription factor (Myt TF) family consists of three zinc finger proteins expressed in neuronal and endocrine cells [including *Myt1 (Nzf2)*, *Myt1l (Myt2, Nzf1)*, and *ST18 (MYT3, Nzf387)*] (15,16). In mice, these factors regulate beta-cell function, survival, and proliferation by repressing several stress-response genes (17–21). In humans, single-nucleotide polymorphisms and deletions in these genes are associated with autism, cancer, craniofacial anomalies, diabetes, or mental illnesses (1,15,22–26). Intriguingly, the protein levels and/or nuclear localization of MYT1 and ST18 were regulated by glucose and free fatty acids in primary human beta-cells, and the reduced MYT TF production preceded beta-cell dysfunction during T2D development (18).

The above studies suggest that the Myt TFs are environment-responsive regulators that prevent mouse beta-cell failure. Our objective here is to investigate this possibility in primary human beta-cells, focusing on *MYT1* and *ST18* because they, but not *MYT1L*, are regulated by high glucose or free fatty acids in human islets (18). For this goal, transplantation of manipulated human islets into mouse models was used to mimic *in vivo* cell responses to obesity-related stress.

## Methods

### Human donor islet and mouse usage

Human donor islets, de-identified (exempt from Institutional Review Board review), were obtained from the IIDP or Dr. A.N. Balamurugan’s islet cell laboratory (University of Louisville, KY, USA) (27), who obtained informed consent before islet isolation. Islets with >80% purity, >95% viability, and a stimulation index>2.0 were used. Mice were used in accordance with protocols approved for Guoqiang Gu by the Vanderbilt University Institutional Animal Care and Use Committee. All policies of the Office of Laboratory Animal Welfare (breeding, surgery, and euthanasia) were followed.

Lentiviral production for gene knockdown in pseudo-islets

Each knockdown (KD) shRNA was expressed in one lentiviral construct driven by a *U6* promoter, co-expressing eGFP for monitoring transfection. For pseudo-islet (PSI) production, dissociated single islet cells were washed with CMRL1066 (10% heat-inactivated FBS, 1X antibiotics, and 1X L-glutamine) and resuspended in the Vanderbilt-Pseudo-islet-media (VPM) (50% CMRL1066 + 50% Vasculife Basal Media, 10% heat-inactivated FBS, 1X antibiotics, 0.5X Glutamax, 2.5 mM HEPES, 0.5X sodium pyruvate plus recombinant human (rh) VEGF, rh EGF, rh FGF basic, rh IGF-1, Ascorbic Acid, Hydrocortisone Hemisuccinate, Heparin Sulfate LifeFactorGentamicin/Amphotericin, and iCell Endothelial Cells Medium Supplement). Cells (200,000 in 200 μL) were aliquoted into one prepared Aggrewells 96-well plate, mixed with 40- 50 ng of lentivirus. The plate was centrifuged at 200g for 5 min and then left undisturbed in a cell culture incubator for 5 days to allow PSI formation. Well preparation uses 100 ul/well of an anti-adhesive rinsing solution (96-well plate, Nacali #4860-900SP) (28).

### Pseudo-islet transplantation and insulin tolerance test

Roughly 30 PSIs were transplanted into anterior eye chambers (AECs) of 2-4 month-old NSG-DTR male or female mice (the Jackson Laboratory, Stock #: 027976), kept in small cages in a pathogen-free facility with a 12-hour dark-light cycle, with free access to food and water (29). Transplants were recovered 4 weeks after transplantation or after another 12 weeks on a control diet (∼13% calories from fat, Lab Diet) or a high-fat diet (HFD, ∼60% calories from fat, Fisher Scientific) challenge. For the insulin tolerance test, mice were fasted for 4 hours, injected with Humalog insulin (0.5 Units/kg) and assayed for blood glucose at 0, 15, 30, 45, and 60 minutes.

### TUNEL assays and Immunofluorescence staining

TUNEL assays used kits from Invitrogen. Immunofluorescence (IF) staining was down on paraffin or frozen sections. Antibodies were: rabbit or guinea pig anti-Pdx1 (1:5000, or 1:2000) (gifts from Chris Wright, Vanderbilt University); goat anti-somatostatin (1:500) (Millipore); rabbit anti-somatostatin (1:2000) (Santa Cruz); rabbit anti-pancreatic polypeptide (1:500) (Abcam); guinea pig anti-insulin (1:1000) and mouse anti-glucagon (1:5000) (Dako); rabbit anti- Myt1 (1:1000) and rat anti-St18 (1:1000) (this lab); rabbit anti-MafA (1:1000, Novus); and chicken anti-Nkx6.1 (1:500) (Developmental Studies Hybridoma Bank, F55A10). Validation used antigen-negative tissues/cells. Secondary antibodies were from Jackson ImmunoResearch (1:1000, Cat# available upon request). Slides were counterstained with DAPI, imaged with laser-scanning confocal microscopy or Airyscan (FV1000 or Leica 880), and quantified with ImageJ.

### Secretion assays

PSIs were pre-incubated in KRB (with 2.8 mM glucose or G2.8) at 37 °C for 1 hour, split into 3 or 4 wells (15-20 PSIs each), and stimulated with G2.8 or G16.7 (16.7 mM glucose) for 45 minutes to quantify secretion. PSIs were lysed with ethanol/HCl to obtain total insulin/glucagon amounts. KCl-induced secretion used 30 mM KCl. Insulin or glucagon quantification was performed using ELISA kits from Alpco or Mercodia, respectively.

### Real-time quantitative PCR (qPCR), scRNA-seq, bulk RNA-seq, and function annotation

RNA was isolated using DNA-free RNA^TM^ kits (Zymo Research). For qPCR, RNA was converted to cDNA (Promega’s High-Capacity cDNA Synthesis Kit) and analyzed with the Bio-Rad SYBR Green Master Mix. Oligos were: *GAPDH*: 5’-ctttggtatcgtggaaggactc-3’ and 5’- agtagaggcagggatgatgt-3’; *MYT1*: 5’-ggccacatcaccgggaacta-3’ and 5’-agtgggcagccatgaggttt-3’; *MYT1L*: 5’-cagctgctgccatcctgaac-3’ and 5’- gcccgctgtttgatggtcag-3’; and *ST18*: 5’- ggcatgcagactctgtggct-3’ and 5’-catccacagccagccattcg-3’.

For RNA-seq, 2-3 technical replicates were included per donor-KD condition. RNA preps with RIN 7.6 were sequenced on the Novaseq 6000, yielding 100-120 million reads per sample. The gene count matrix (60,683 total genes) was obtained using Genialis (30) and lowly expressed genes were excluded by requiring a minimum of 3 reads in at least 3 of the 12 samples, resulting in 34,266 genes retained for downstream analysis (ESM Table 1). Differentially expressed genes (DEGs) were identified using DESeq2 (31) with a paired design, and *p*-values were adjusted for multiple testing using the Benjamini–Hochberg procedure applied to all 34,266 expressed genes, a method suitable for small sample sizes. DEGs were defined by a log2 fold change threshold of 0.5 and an FDR-adjusted *p*-value cutoff of 0.05. Functional enrichment analysis was performed against Gene Ontology (GO) database with ClusterProfiler (32). Additionally, gene set enrichment analysis (GSEA) was conducted using the fgsea R package with genesets (35,134 total) from MSigDB (version 2025.1.Hs).

For scRNA-seq, dissociated PSIs with ∼80-90% single cells and <5% dead cells were used for InDrop, sequenced with Novaseq 6000 (Illumina) with ∼120 million reads per sample. DropEst was used to generate count matrices. Cells with low unique mapping reads (<500), low proportion of expressed genes (<100), or high proportion of mitochondrial RNAs (>10%) will be excluded (4). Reads were normalized using UMI-filtered counts. Cell subpopulations were identified and visualized using UMAP in Seurat, based on the first 30 principal components generated from the top 2000 highly variable genes (33,34). Seurat identified DEGs at the criteria of |log2 fold-change|> 0.25 & FDR≤ 0.05. Pathway analysis used Database for Annotation, Visualization and Integrated Discovery (DAVID) for functional clustering (35) (*p*adj < 0.05, Benjamini-Hochberg procedure), enabling functional clustering of similar processes for data interpretation. To identify the specific functions, each gene was searched for association with secretion, stress, or cell death/apoptosis using the gene STRING (36). We then verified the annotation by examining published data.

### Ca^2+^ imaging

Dissociated islet cells attached to glass-bottomed plates (D35-14-1.5P, Cellvis) were infected with shRNA-expressing lentivirus for 3 days, and with RCaPMP-expressing adenovirus (beta-cell specific) overnight. Cells were then incubated (20 minutes, 37°C) in KRB with 1 mM glucose (G1). Real-time fluorescence imaging (5 seconds per frame) followed with a 20X objective, recordings for 5 minutes under G1, 25 minutes under G11, and 5 minutes under 30 mM KCl.

### Overlapping MYT-regulated and T2D-associated genes

For PSI- or beta cell-specific gene overlapping assays by hypergeometric tests, we used the 34,266 PSI (ESM Table 1) or 18,752 beta cell genes [ESM Table 2, expressed in beta cells with TPM > 0.5 (18,350) in (37) plus those extras from Walker et al. (38) or our scRNA-seq], as starting populations, respectively. Six T2D gene lists were used: all the T2D-associated genes that are expressed in PSIs (2,047 genes, ESM Table 3) or beta cells (1,806, ESM Table 4) from the GWAS catalog [p<9X10^-6^, (39)]; T2D genes expressed in human PSIs (665, ESM Table 5) or beta cells (621, ESM Table 6) from the Suzuki et al T2D-risk gene studies (p<5X10^-8^) (1); and the DEGs between healthy and T2D donor islets (686, ESM Table 7) or beta cells (365, ESM Table 8) of the Walker studies (38).

### Statistical analysis

Paired t-tests, repeated-measures ANOVA, or linear mixed-effects models were used for pairwise comparisons at single time points/paired genotypes or across multiple groups of data points. An adjusted p-value £0.010 (for GSEA) or ≤ 0.050 (all else) was considered statistically significant.

### Data accessibility and request for materials

The RNA-seq data were deposited in the Gene Expression Omnibus (GEO) (GSE307815 for scRNA-seq and GSE307751 for bulk RNA-seq). Guoqiang Gu is the guarantor of the data, and he will fulfill reasonable resource and reagent requests.

## Results

### High levels of ST18 are required for human beta-cell GSIS

Using lentivirus-driven shRNAs identified via reporter assays (ESM Fig. 1), we reduced the MYT1 or ST18 mRNA levels by (79.9 ± 2.5 )% (mean ± SEM) or (74.2 ± 8.3)%, respectively, in human PSIs (Fig. 1a-d). MYT1-KD did not change the % of insulin secreted or stimulation index (secretion at G16.7/that at G2.8) under 16.7 mM glucose (G16.7) (Fig. 1e and f). In contrast, ST18-KD resulted in a statistically significant reduction in GSIS compared to controls, both the % of total insulin secreted [(52.0 ± 9.1)% reduction] (Fig. 1e) and the stimulation index (26.7 ± 7.2)% reduction] (Fig. 1f).

**Fig 1.**
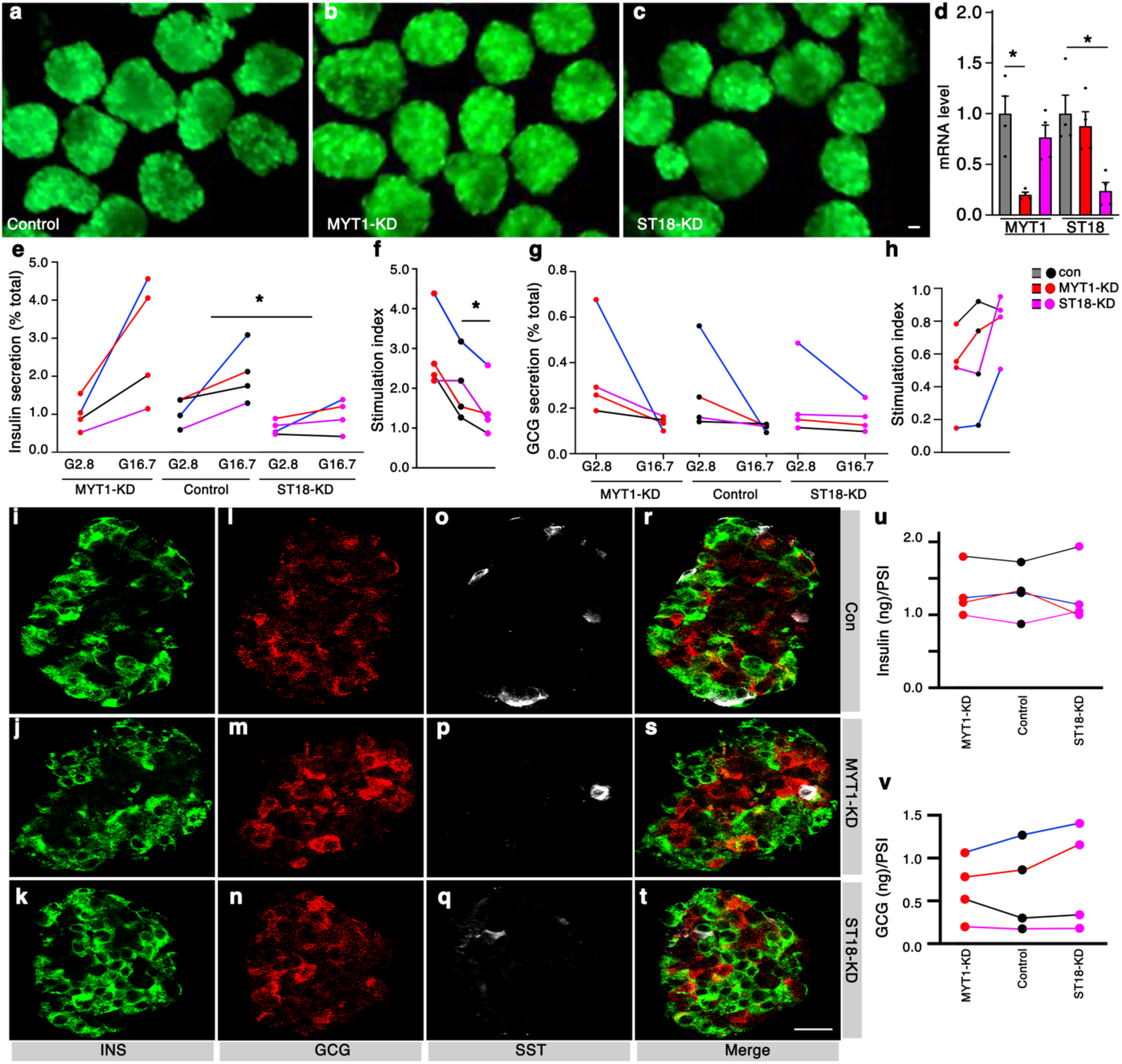
*ST18*-KD, but not *MYT1*-KD, impairs GSIS of primary human beta cells. Human islets, after quality check, were dissociated into single cells, infected with an eGFP-expressing lentivirus that also expressed control, *MYT1*-targeting, or *ST18*-targeting shRNAs, and then used to make PSIs and tested. (a-c) Representative PSI images. (d) RT-PCR assays showing *MYT1*- or *ST18*-KD levels in PSIs made from four donors. The grey bars in the *MYT1* or *ST18* assay subgroups were shared controls for the *MYT1*- or *ST18*-KD samples. Each dot represented data from one donor. Shown were mean + SEM. *: *p*<0.05, from t-test (two-tailed type two errors). (e-h) PSI secretion of insulin (e and f) or glucagon (GCG) (g and h) in response to G2.8 or G16.7, presented as % of total hormone secreted (e and g) or stimulation index (f and h). Connected lines showed islets from the same donor. Each dot represented the average of 3-4 technical replicates from the PSIs of one donor. The color of the connecting line indicated donor identity (black, blue, red, pink lines corresponded to donor #1, #2, #3, and #4, respectively). *P* values in e were from repeated measure ANOVA, assaying the combined effect of gene-KD and glucose stimulation. *P*-values in f were from a paired t-test (two-tailed type 2 errors). *: *p*<0.05. (i-t) Hormone expression in newly produced PSIs, represented by samples of donor #1. Single channels (i-q) and merged panels (r-t) were shown. (u and v) Insulin and glucagon content in PSIs after *MYT1*- or *ST18*-KD, normalized against the numbers of PSIs of similar sizes. Each dot in u and v represented the result of one donor of 3-4 technical replicates. Connected dots were from the same donor. The color of connecting lines indicated donor identity (black, blue, red, pink lines corresponded to donor #1, #2, #3, and #4, respectively) Scale bars, 20 μm.

There was no difference in glucagon secretion in control, MYT1-KD, and ST18-KD PSIs (Fig. 1g and h). MYT1- or ST18-KD did not cause hormone co-expression between insulin, glucagon, and somatostatin (Fig. 1i-t) or insulin, pancreatic polypeptide, and somatostatin (ESM Fig. 2a-l). The levels of insulin or glucagon were not significantly affected by KD (Fig. 1u and v), neither TFs such as PDX1, MAFA, and NKX6.1 (ESM Fig. 2m-u). These findings suggest that high MYT1 or ST18 levels are dispensable for beta-cell identity, while ST18 is required for GSIS.

**Fig 2.**
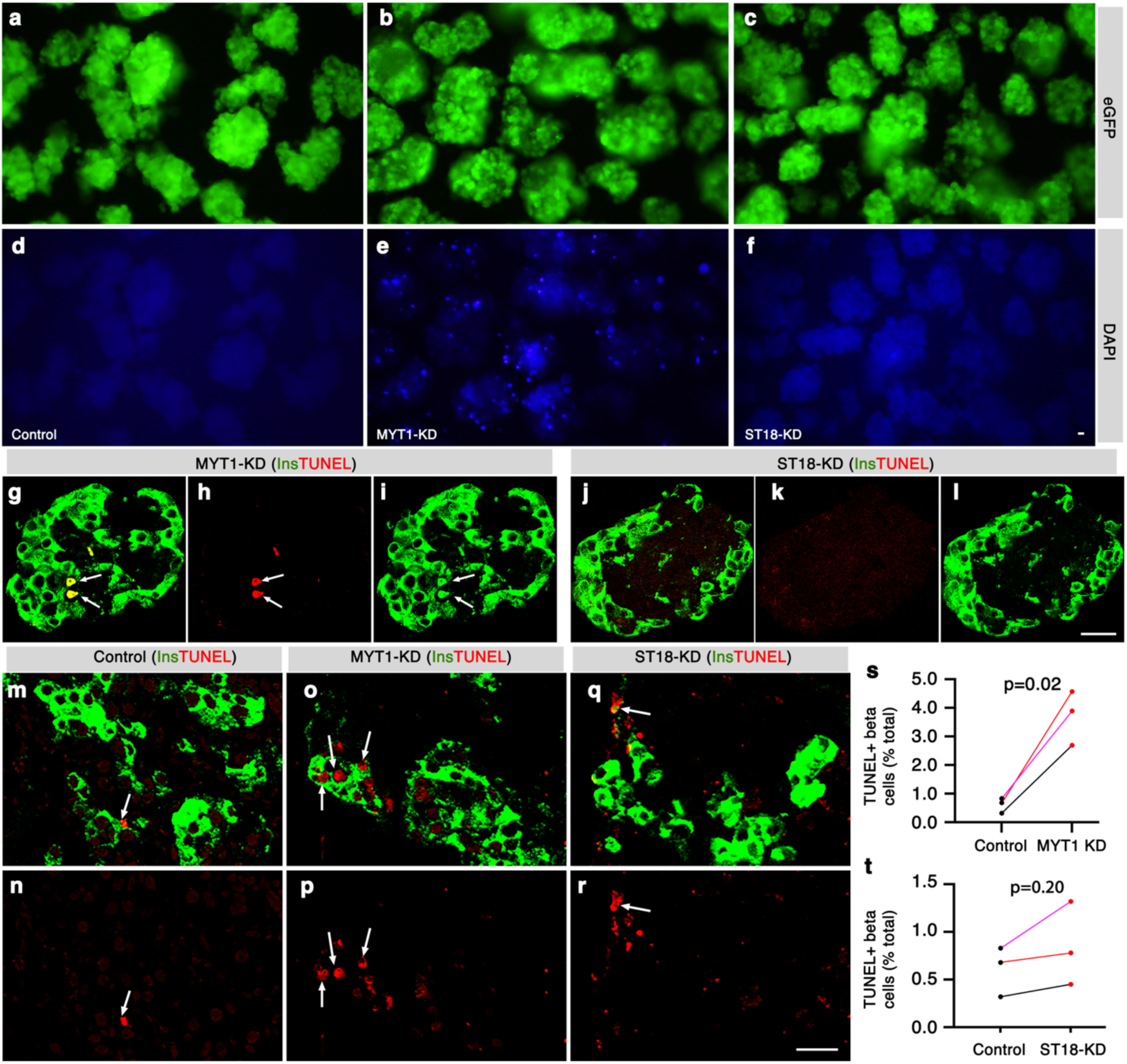
*MYT1*-KD, but not *ST18*-KD, causes the death of human beta cells *in vitro* and *in vivo* under normal physiology. Freshly made PSIs, with control, *MYT1*-KD, or *ST18*-KD, were stained with DAPI to visualize dead cells. They were also examined for cell death with TUNEL assays before or 4 weeks after transplantation. (a-c) PSIs with control, *MYT1*-, or *ST18*-targeting shRNA expression (with green eGFP reporter). (d-f) The corresponding DAPI in panels a-c. (g-l) TUNEL staining of *MYT1*-KD (g-i) and *ST18*-KD (j-l) PSIs. White arrows pointed at two dead cells with Insulin-signals. (m-t) TUNEL assays and quantification of dead beta cells 4 weeks after PSIs were transplanted into AEC of mice. Merged (m, o, p) and a single TUNEL channel (n, p, r) for each condition were shown. White arrows pointed at several dead insulin cells. In s and t, each dot represented results from one donor, transplanted into at least three mice. *P* values were from paired t-tests (with data from the same donor pair). Connecting lines indicated islets from the same donor. The color of the connecting line indicated donor identity (black, red, pink lines corresponded to donor #1, #2, and #4, respectively).*: *p*<0.05. Bars, 20 μm.

### High levels of MYT1 are required for beta-cell survival under normal physiology

Freshly prepared PSIs were stained briefly with DAPI^+^ to label the nuclei of dead cells. PSIs with MYT1-KD but not ST18-KD contained DAPI cells (Fig. 2a-f). TUNEL assays detected DNA fragmentation in MYT1-KD beta cells, suggesting their apoptosis (Fig. 2g-l).

Because human beta-cell death detected in vitro was sometimes not shown in vivo (40), we examined how beta cells with MYT1- or ST18-KD behaved as xenotransplants in mice. PSIs were transplanted into the AECs of NSG-DTR mice, followed by cell death assays four weeks later. In all three donor samples tested, we found statistically more [(6.6 ± 1.07)-fold] beta-cell death upon MYT1-KD but not ST18-KD compared to controls (Fig. 2m-t and ESM Fig. 3a-c), corroborating the in vitro data. Note that islets from one donor (#3, ESM Human-Islet-Checklist) had a low yield of PSIs and were not tested for in vivo cell death.

**Fig 3.**
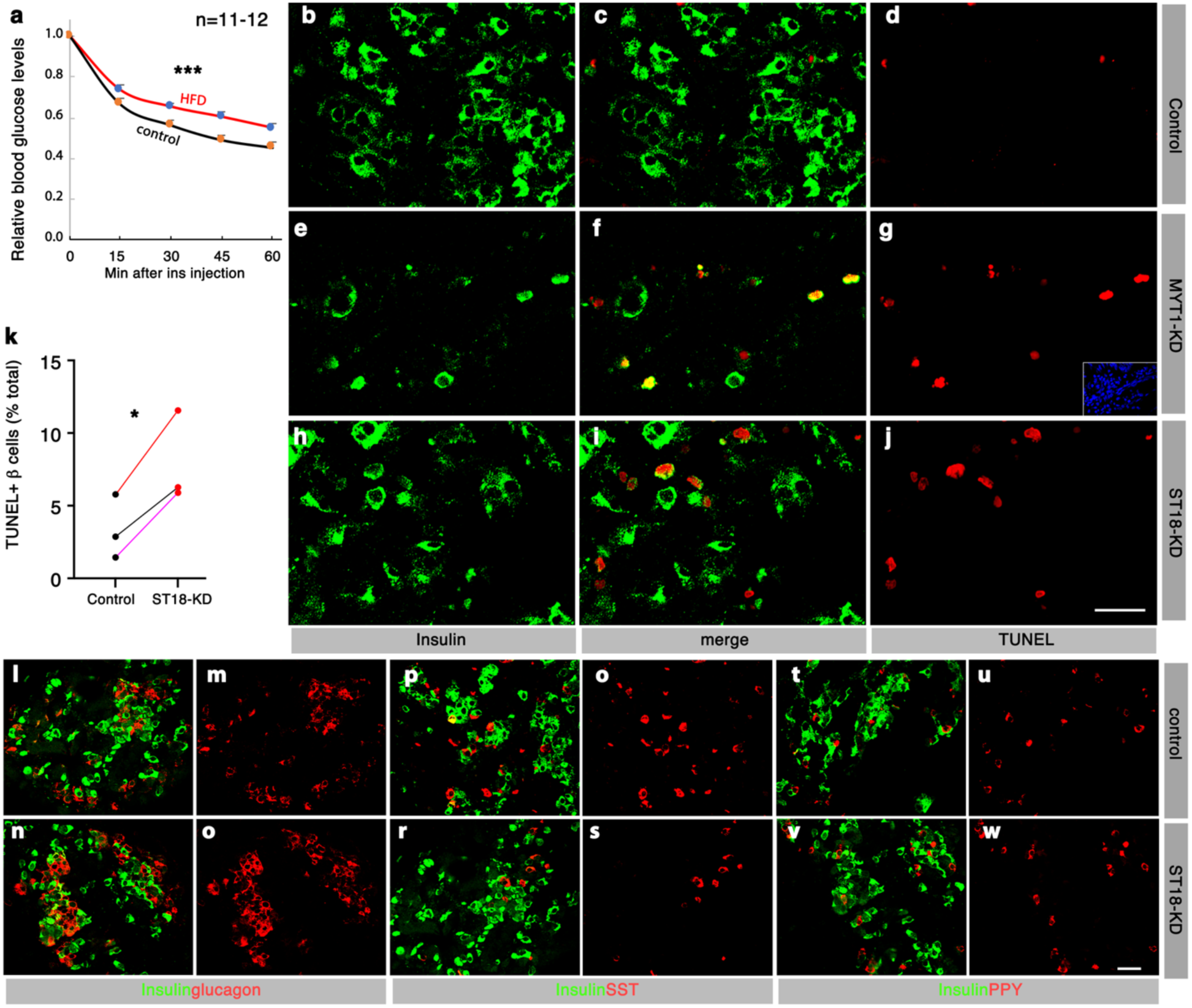
*ST18*-KD renders human beta cells vulnerable to HFD-induced death. Four weeks after the operation, mice transplanted with manipulated PSIs were fed with HFD for 12 more weeks. PSIs were then recovered for characterization. (a) HFD-induced insulin resistance in recipient mice. The starting blood glucose of each animal was normalized to “1”. The *p*-value is from a linear mixed-effects model. ***: *p*<0.001. (b-k) TUNEL assays of recovered transplants. Single insulin (b, e, h) and TUNEL (d, g, j) and merged panels (c, f, i) were shown. Inset in g showed a DAPI staining in processed sections to visualize all nuclei. In k, each dot represented data from one donor, including at least three transplanted mice. Connecting lines indicated islets from the same donor. The color of the connecting line indicated donor identity (black, red, pink lines corresponded to donor #1, #2, and #4, respectively). “*p*” is from paired t-tests. *: *p*<0.05. (l-w) IF showing the hormone co-expression status between insulin and other hormones in recovered transplants, with merged images (l, n, p, r, t, v) and single glucagon (m, o), somatostatin (SST) (q, s), and pancreatic polypeptide (PPY) (u, w) channels shown. Bars, 20 μm.

### *ST18*-KD renders beta cells vulnerable to obesity-induced death

We examined how MYT1- or ST18-KD affected the response of human beta cells to obesity. Mice with xenotransplants were fed with an HFD for 12 weeks, starting 4 weeks after transplantation. This did not induce overt diabetes but induced insulin resistance (Fig. 3a). Consistent with the death of MYT1-KD beta cells, very few insulin^+^ cells were detected in MYT1- KD transplants (Fig. 3b-g and ESM Fig. 3d-f). More beta-cell death [(2.7 ± 0.7)-fold] was detected in transplants with ST18-KD than in controls (Fig. 3h-k and ESM Fig. 3d-f). We did not detect insulin^+^ cells activating other islet hormones (Fig. 3l-w). In addition, beta-cell death was not observed in ST18-KD samples without HFD challenge after prolonged transplantation (ESM Fig. 3g and h). These findings suggest that ST18 is not required for beta-cell identity but needed for beta-cell survival under obesity-related stress.

### *MYT1*-KD or *ST18*-KD deregulates genes involved in cell death or secretion

Out of the 34,266 expressed genes in PSIs (ESM Table 1), there are 495, 79, and 735 DEGs identified in paired comparisons between MYT1-KD versus control, ST18-KD versus control, and ST18-KD versus MYT1-KD samples, respectively (Fig. 4a, ESM Fig. 4a-c and ESM Tables 9-11). Based on the GO enrichment analysis, the DEGs from MYT1-KD versus control are enriched for organelle membrane, granule, protein catabolism, cell junction, ion transport, etc. (ESM Table 12a). Microtubule binding, shown to regulate insulin secretion (41,42), was enriched in the DEGs from ST18-KD versus control (ESM Table 12b). Similarly, the DEGs of ST18- KD versus MYT1-KD are enriched for those regulating membrane, secretion, death, etc. (ESM Table 12c).

**Fig. 4.**
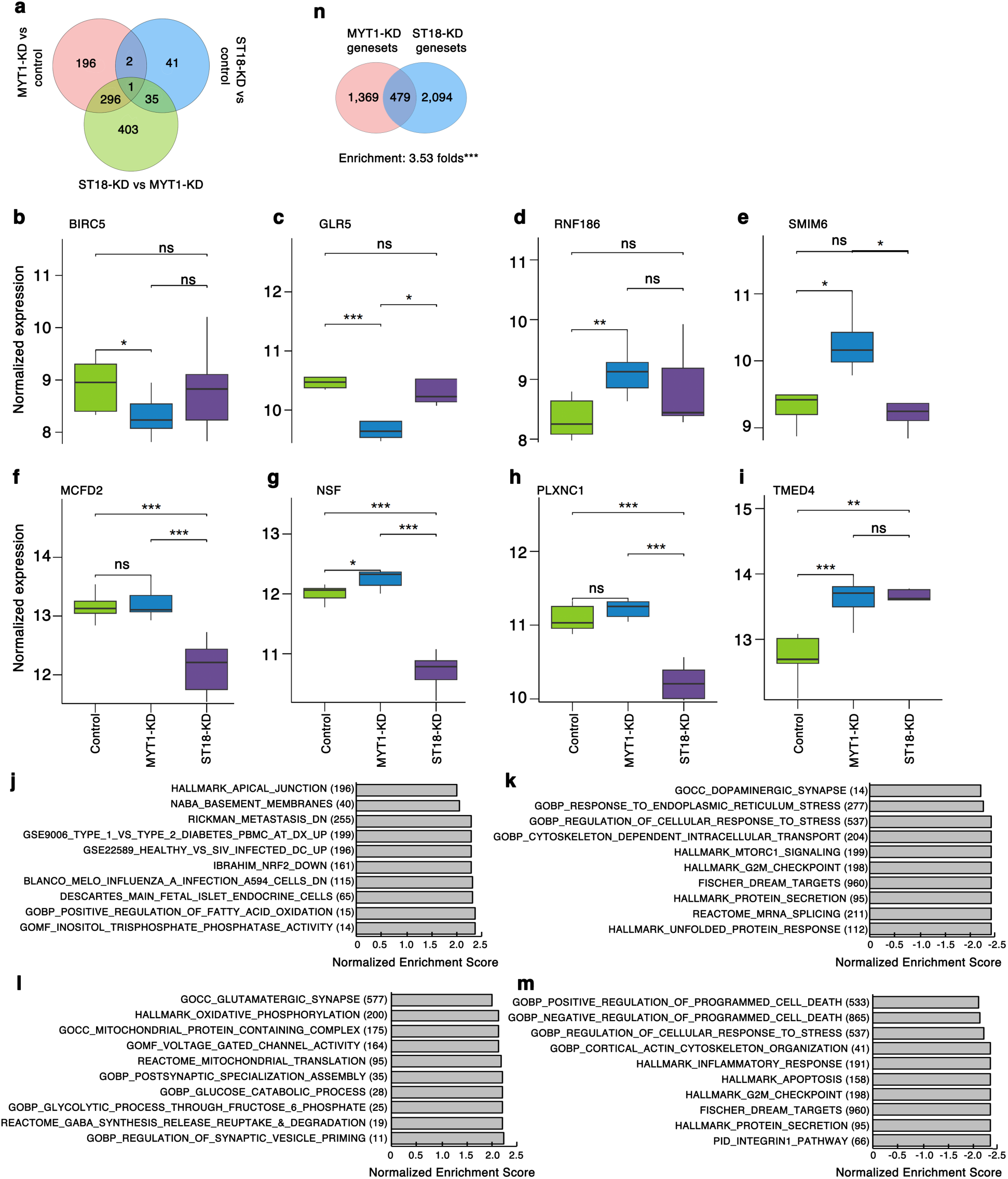
Human PSIs with *MYT1*- or *ST18*-KD deregulate overlapping but distinct genes/genesets, especially those involved in cell death and secretion. DESeq2 and GSEA were used to identify DEGs or dysregulated pathways. (a) A Venn diagram showing the number and overlapping status of DEGs when comparing the expression between samples of: *MYT1*-KD vs. controls, *ST18*-KD vs. controls, and *ST18*-KD vs. *MYT1*-KD. The single gene shared by all three DEG list is *CEND1*, important for neuronal differentiation but its function in beta cells is unknown. (b-i) Expression of a few selected DEGs involved in cell death/apoptosis, stress response, and secretion related genes in control, *MYT1*-KD, and *ST18*-KD PSIs. Y axis shows the normalized expression values after variance stabilizing transformation (VST) in DESeq2. Shown are results (box and whisker plot) from 4 biological samples (2 or 3 biological replicates per sample). (j and k) Selected up- or down-regulated pathways in *MYT1*-KD PSIs vs controls. (l and m) Selected up- or down-regulated pathways in *ST18*-KD PSIs vs controls. For each geneset in j-m, the numbers in parentheses indicate the number of leading-edge genes in that geneset. (n) Number of overlapping pathways between those affected by *MYT1*- or *ST18*-KD compared to controls (***: p<0.001, hypergeometric analysis).

We inspected the DEGs individually, focusing on those regulating cell death/apoptosis, stress response, and secretion. Thirty-three of the 495 DEGs from MYT1-KD versus control have established roles in these processes (ESM Table 9). Underscoring the complexity of these processes, we found both upregulated and downregulated cell death/stress-response genes, including both positive and native regulators. For example, BIRC5 and GLR5 (both inhibit cell death in response to increased cellular or excitatory stress, see ESM Table 9 for references) were downregulated in MYT1-KD PSIs (Fig. 4b and c). In contrast, RNF186 and SMIM6, with RNF186 inducing cell death in response to ER stress while SMIM6 inhibiting it, were both upregulated (Fig. 4d and e). From the 79 ST18-KD DEGs, nine were involved in cell death, stress response, or secretion (ESM Table 10). For example, MCFD2, NSF, and PLXNC1, all promoting secretion, were downregulated (Fig. 4f-h), while TMED4, inhibiting ER stress, was upregulated (Fig. 4i). These findings are consistent with the reduced cell viability or GSIS in MYT1- or ST18- KD PSIs, respectively.

### MYT1 and ST18 regulate common and unique genesets in PSIs

GSEA showed that MYT1-KD, compared to controls, deregulated 1,848 genesets (ESM Table 13). The upregulated genesets regulate glucose metabolism, mitochondria, channels, gamma-aminobutyric acid (GABA)-regulated secretion, etc. (Fig. 4j), which either promote GSIS [e.g., glucose metabolism (43)] or inhibit it [e.g., GABA_synthesis (44)]. The downregulated genesets regulate stress response, apoptosis, etc. (Fig. 4k). Similarly, ST18-KD in PSIs significantly deregulated 2,573 genesets (i.e., pathways) over controls (ESM Table 14), including upregulated pathways in fatty acid oxidation, oxidative stress (NRF2-related), etc. (Fig. 4l), and downregulated pathways in ER stress, protein secretion, etc. (Fig. 4m). Consistent with the overlapping and different activities of the MYT TFs, MYT1-KD and ST18-KD shared 479 deregulated genesets (ESM Table 14), a statistically significant 3.53-fold enrichment (p<0.001, Fig. 4n); there were also 1,379 differentially expressed genesets between the MYT1-KD and ST18-KD PSIs (ESM Table 15, ESM Fig. 4d and e). Also consistent with the roles of MYT TFs in stress response, both MYT1-KD or ST18-KD PSIs have deregulated the “Regulation_of_cellular_response_to_stress” pathways (Fig. 4k and m).

### ScRNA-seq revealed ST18-dependent genes in beta cells

High-quality sequencing data were obtained from one batch of PSIs, detecting all the major islet cell types and non-islet cells in PSIs (alpha, beta, delta, and gamma) (Fig. 5a) (ESM Fig. 5). These contained a total of 536, 906, and 661 beta cells in control, MYT1-KD, and ST18- KD samples, respectively. There was no clear separation between control and MYT1-KD beta cells (Fig. 5b); neither difference in their MYT1 expression, despite the presence of 687 DEGs between them (ESM Table 16). These results suggest the MYT1-KD scRNA-seq data are mostly from cells with minimal MYT1-KD due to the death of MYT1-KD beta cells [which could affect live beta cells non-cell autonomously (45)]. These MYT1-KD data were excluded from the current report.

**Fig. 5.**
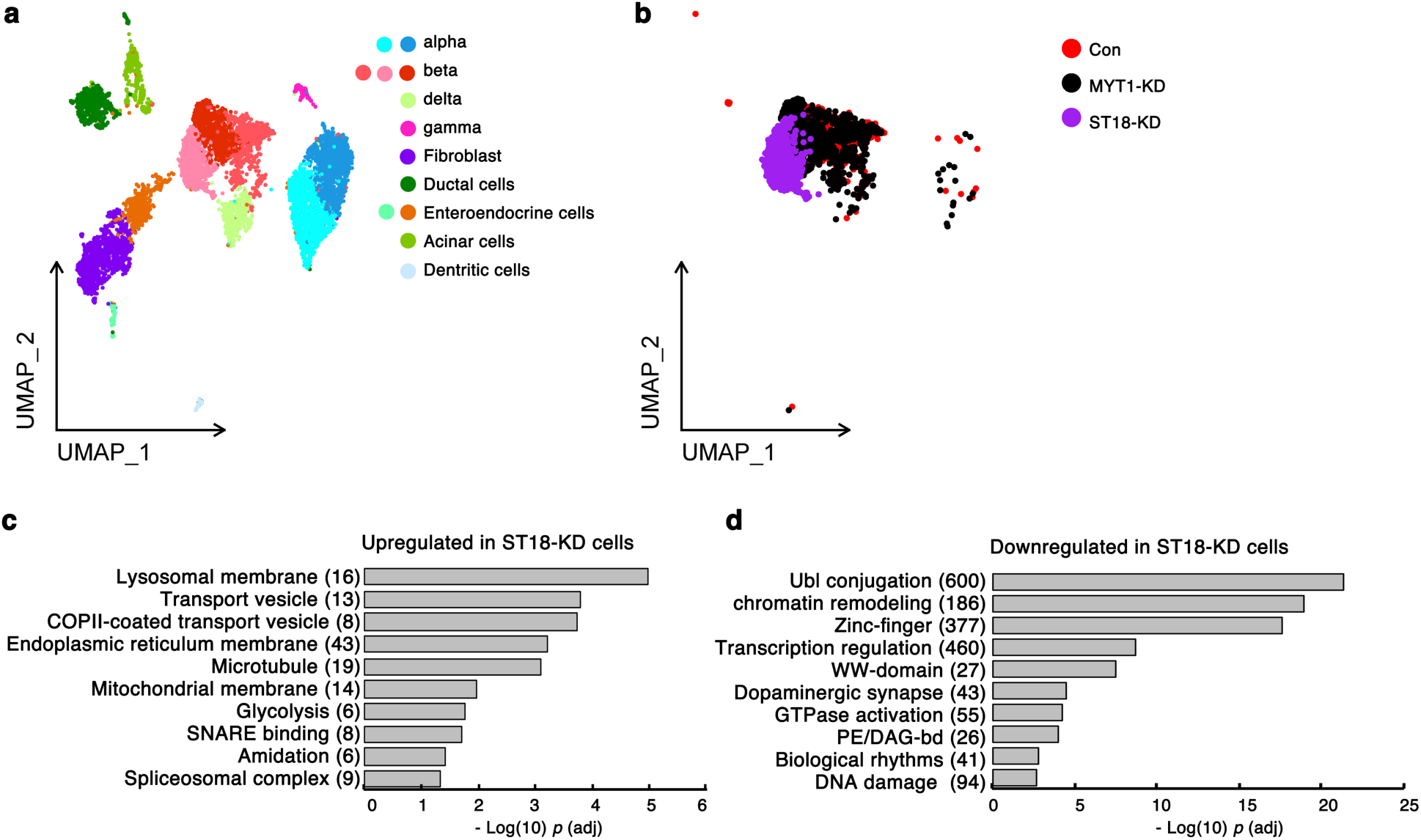
*ST18*-KD deregulates several processes involved in beta-cell secretory functions. Freshly prepared PSIs were dissociated and used for InDrop scRNA-seq. (a and b) UMAPs showing the separation of cells in PSIs, including all cells (a) or beta cells only (b). (c and d) Pathways deregulated in *ST18*-KD human beta cells, composed of up- (c) or down-regulated (d) genes in mutants. The number in parentheses following each process refers to the number of genes discovered. *P*-values were adjusted with the Benjamini-Hochberg procedure.

The control and ST18-KD beta cells were clearly separated and had 3,284 DEGs, including ST18 (Fig. 5b, ESM Table 17). ST18-KD upregulated several membrane-related processes, SNARE binding, and mitochondrial membrane (Fig. 5c) while down-regulating Ubl conjugation, biological rhythms, etc. (Fig. 5d). Consistent with the death of ST18-KD beta cells in vivo under stress, HSPA1A, whose overexpression led to mouse beta-cell death in vivo (18), was activated in ST18-KD beta cells (ESM Tables 17). These results led us to probe how ST18 regulates GSIS, focusing on mitochondria- and Ca^2+^ influx-related processes.

### *ST18*-KD compromises glucose-stimulated Ca^2+^ influx in beta cells

Consistent with the upregulated mitochondrial membrane proteins (Fig. 5c), the ST18- KD beta cells had higher mitochondrial volume (20.1 ± 2.2%) (ESM Fig. 6a-e). But this did not result in higher mitochondrial transmembrane potential in mutant cells (ESM Fig. 6f). The numbers of insulin granules in control and ST18-KD beta cells were similar (ESM Fig. 6g-k).

**Fig. 6.**
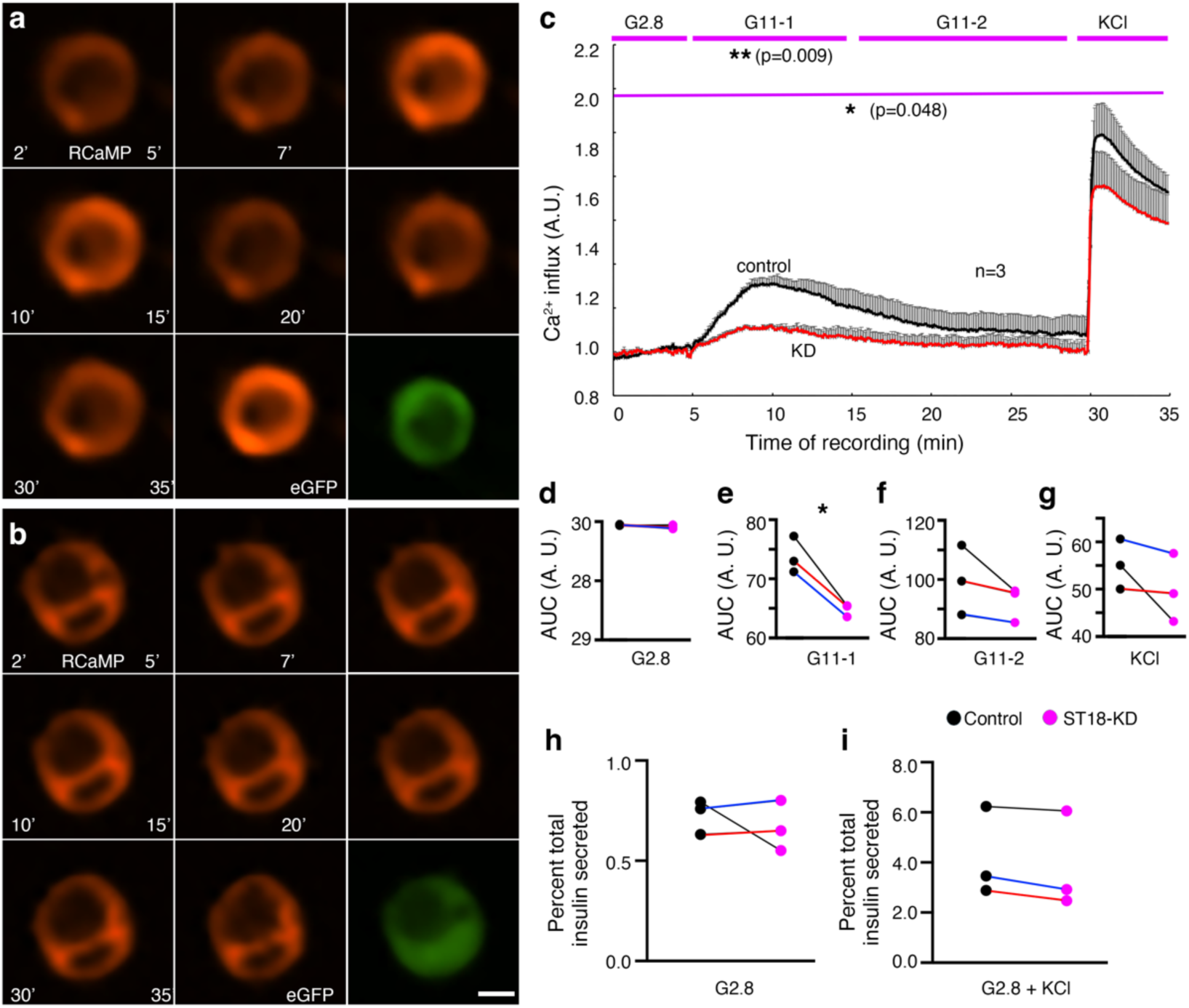
*ST18*-KD compromises glucose-induced Ca^2+^ influx in beta cells. (a-g) The effect of *ST18*-KD on Ca^2+^ influx in beta cells, shown by the intensity change of RCaMP fluorescence over time after 11 mM glucose or 30 mM KCl addition (at 5-minute or 30-minute mark, respectively). One control (a) and one *AT18*-KD cell (b) were shown. eGFP-expression (green, the last panel in each group) was used to identify cells with control or targeting shRNA expression. Bar, 5 μm. (c) RCaMP fluorescence intensity changes over time in beta cells of three donors. 126, 138, 127 control cells (from donor #5, 6, and 7, respectively) and 138, 102, and 123 *ST18*-KD cells (from donor #5, 6, and 7, respectively) were assayed. The results from each donor’s cells were averaged and treated as one sample (represented by one dot from d to g). The (mean + SEM) values were presented. *P* values were from repeated measure ANOVA, analyzed at separate phases of the recording [G2.8, 10 minutes of G11 (G11-1), 15 more minutes G11 (G11-2), and 5 more minutes of 30 mM KCl] or the entire process (**: p<0.01; *: *p*<0.05). The relative area-under-curve (AUC) for each phase was also presented (d-g. *: *p*<0.05, t-test with two-tailed type two error). (h and i) Insulin secretion in control and *ST18-KD* PSIs in response to KCl-induced depolarization in 45-min windows. Each dot represented the average of 3-4 technical replicates from one donor. The connected dots indicated results from the same donor. The colored connecting lines in d-j indicated the donor identity (black: donor #5, red: donor #6, blue: donor #7).

A down-regulated process in the ST18-KD beta cells was “dopaminergic synapse” (Fig. 5d), including voltage-gated Ca^2+^ channels CACNA1A, CACNA1B, CACNA1C, and CACNA1D (ESM Table 17). In addition, several K^+^ and Na^+^ channel genes were affected by ST18-KD (ESM Table 17). Consequently, the ST18-KD compromised high glucose-induced Ca^2+^ influx, with the most statistically significant effect occurring within the first 10 minutes of glucose stimulation (Fig. 6a-g). In addition, ST18-KD beta cells had statistically indistinguishable KCl-induced Ca^2+^ influx (Fig. 6c and g) and insulin secretion (Fig. 6h and i). These results are consistent with the conclusion that the defective glucose-induced Ca^2+^ influx, a major cause of the reduced GSIS in ST18-KD beta cells, is likely a combined result of deregulated Ca^2+^, K^+^, Na^+^ channels, and other processes. Each of these factors may have a weak effect on Ca^2+^ influx when examined individually, but their combined effect is statistically significant. Note that ST18-KD beta cells upregulated several SNARE binding protein genes that usually promote GSIS (NSF, SYT5, SYT7, STX1A, STXBP1, and VAMP2) (ESM Table 17). It is likely that these upregulated genes cannot fully compensate for the reduced Ca^2+^ influx to maintain GSIS, or that these SNARE components are not the secretion-limiting factors in this setting.

### Human *MYT1*- and *ST18*-regulated genes are enriched for T2D-associated genes

We tested whether MYT1/ST18 regulated T2D risk genes. We used several published data sets: 1) the 3,319 T2D-risk genes in GWAS Catalog (39), including 2,047 PSI- and 1,806 beta cell-expressed genes (ESM Table 3 and 4); 2) the 849 T2D-associated genes of Suzuki et al. (1), including 665 PSI- (ESM Table 5) and 621 beta cell-expressed genes (ESM Table 6); 3) the 709 islets (686 expressed in PSIs, ESM Table 7) or 365 beta cells DEGs (ESM Table 8) between control and T2D donar islets.

For MYT1-regulated genes, we used the 495 DEGs between control and MYT1-KD PSIs. We found 45, 10, or 17 genes that overlapped between these DEGs and the GWAS, Suzuki, or Walker PSI lists. These representing 1.52 (significant, p=0.011), 1.04 (non-significant), or 1.72 (significant, p=0.023) folds enrichment over random gene distribution (hypergeometric tests) (Fig. 7a-c). For ST18-regulated genes, we used beta-cell specific DEGs, which shared 568, 223, or 84 genes with the beta cell-expressed GWAS-, Suzuki-, or Walker-list, all representing statistically significant enrichments (≥ 1.31-fold enrichment, p≤0.004, hypergeometric tests) (Fig. 7d-f).

**Fig. 7.**
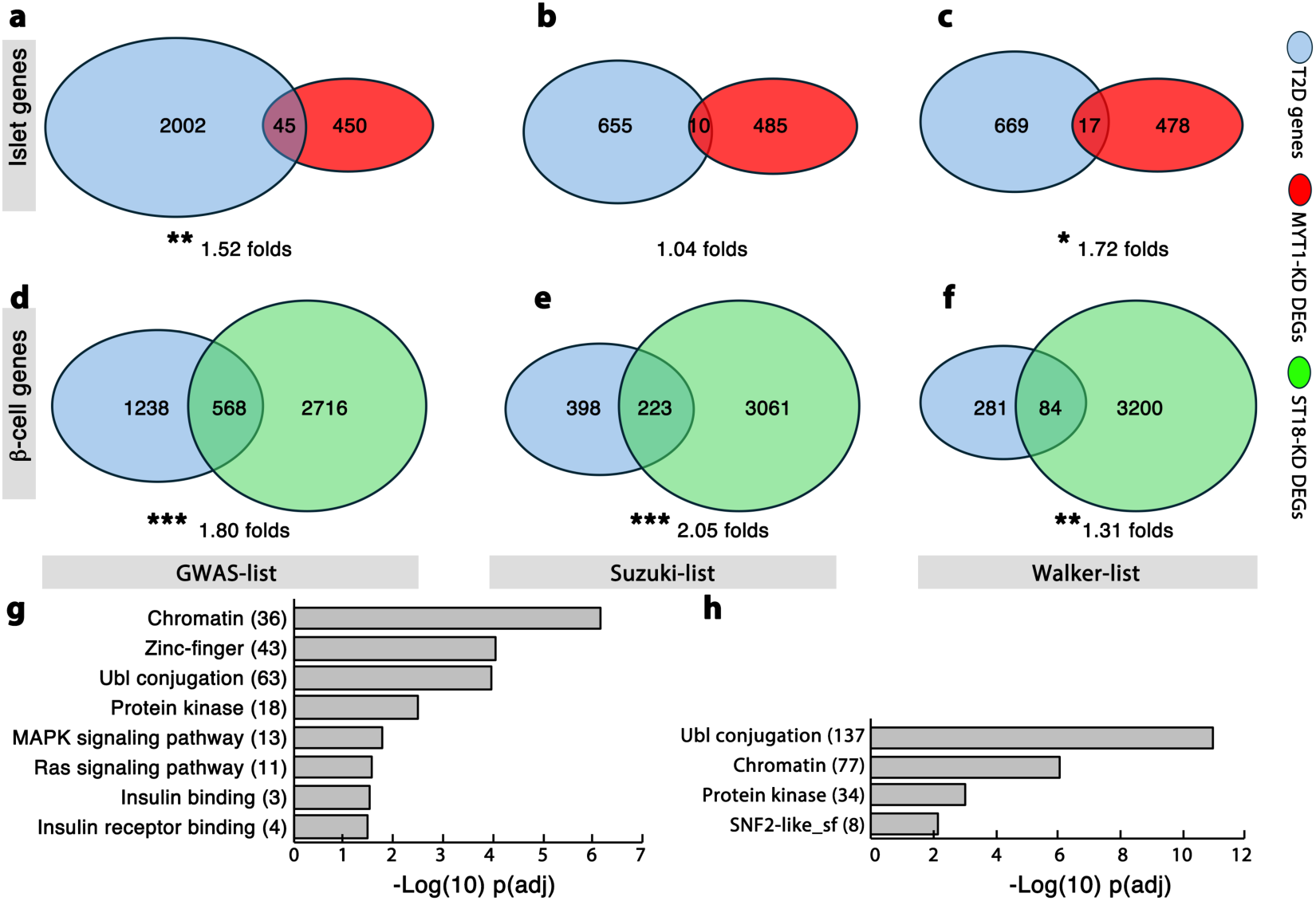
DEGs induced by *MYT1*- or *ST18*-KD are enriched for T2D-associated genes. (a-c) Diagrams showing the number of overlapping genes between the T2D-associated genes and the DEGs induced by *MYT1*-KD in PSIs. All genes with detectable expression in PSIs (ESM Table 1) were used as the starting population in overlapping studies. The enrichment fold and *p*-values were from hypergeometric tests. In (a-c), the GWAS gene list (ESM Table 3), the Suzuki gene list (ESM Table 5), and the Walker gene list (ESM Table 7) were used, separately. (d-f) Similar as a-c, with the DEGs from ST18-KD vs control beta cells and the T2D-associated genes only detected in beta cells from three separate studies (ESM Tables 4, 6, and 8). *: *p*<0.05; **: *p*<0.01; ***: *p*<0.001. (g) Biological processes enriched in the shared genes between T2D-associated beta-cell genes (GWAS list) and *ST18*-KD-induced DEGs in beta cells from scRNA-seq data. (h) Same as those in g, except that the T2D-associated genes were from the Suzuki studies. In g and h, *p*-values were adjusted with the Benjamini-Hochberg procedure. In g and h, the numbers in the parathesis following each pathway indicate the number of genes discovered in that pathway.

We last examined the pathways shared by T2D-risk and ST18-regulated beta-cell genes. Among those shared by the GWAS list and ST18-KD DEGs, chromatin, zinc fingers, Ubl-conjugation, several kinase signaling, and insulin binding processes were enriched (p adj<0.05) (Fig. 7g, ESM Table 17). Among those shared by the Suzuki list and the ST18-KD DEGs, Ubl-conjugation, chromatin, kinase, and SNF2-like processes were enriched (Fig. 7h). These combined results suggest that these MYT TF-regulated processes play key roles in preventing beta-cell failure and T2D.

## Discussion

In this study, we showed that MYT1-KD induced human beta-cell apoptosis while ST18-KD compromised GSIS under normal physiology. ST18-KD also promoted obesity-induced beta-cell death. Corresponding to these cellular functions, MYT1 and ST18 regulated common and unique genes that were enriched for those associated with T2D risk.

Our overall findings suggest that the MYT TFs integrate genetic and nutritional factors to prevent beta-cell failure and T2D. Obesity-associated high glucose or free fatty acids lead to overproduction of unfolded proteins or reactive oxygen species in beta cells. For homeostasis, beta cells activate stress response (8,46). Yet over-activating this response also leads to beta-cell failure. We have reported that in mice, this latter scenario is safeguarded by the Myt TFs (18). We now showed that these MYT-TF functions could be extended to human beta cells, likely involving a few T2D-associated genes and others that regulate beta-cell death and/or functions (47). These new results, together with the regulation of MYT1 and ST18 by nutrients in human beta cells (18), the association of the MYT genes with T2D (39), and the reduced expression of MYT TFs in T2D beta cells (18), highlight MYT TFs as key tunable repressors of obesity-induced beta-cell failure. Manipulating MYT1 and ST18 activities under metabolic stress may be explored to prevent this failure and the development of T2D.

Our studies revealed the different and complementary functions of MYT1 and ST18 (48,49). MYT1 is essential for beta-cell survival, whereas ST18 is essential for high levels of beta cell GSIS, under normal physiological conditions. Yet ST18 also has pro-survival activity under metabolic stress. The unique and overlapping pathways regulated by these factors corroborate these observations. These combined findings highlight a mechanism that multiple processes could regulate a same cellular function under different physiologies, so that different phenotypes would prevail under different conditions when a specific process was perturbed. Thus, testing the functions of specific genesets under different metabolic conditions may reveal nuanced mechanisms of beta-cell protection in obesity.

Our findings also highlighted the species-specific roles of the MYT TFs, despite their high levels of sequence conservation (18). MYT1-KD caused human beta-cell death, while Myt1 inactivation compromised mouse beta-cell function but not survival (21). Similarly, ST18-KD results in human beta-cell death under obesity, yet a mouse St18 protein level reduction did not (50). Furthermore, mouse beta cells with Myt1-single and Myt TF-triple mutations had compromised identity under metabolic stress, but MYT1- or ST18-KD human beta cells had not. These findings highlight the species-specific functions of MYT-TF in mouse and human beta-cell identity maintenance .

Several issues remain to be addressed. First, gene KD may not fully reveal the molecular changes underlying beta-cell dysfunction or apoptosis due to donor differences, varying gene KD levels with shRNA, and the limited sample size (n=4). Thus, the identified DEGs and differentially expressed genesets are considered preliminary, and correlating gene expression changes with cellular defects requires caution. Second, how MYT1 regulates beta-cell viability remains incompletely understood. We detected significantly reduced MYT1 and altered expression of several cell-death regulators in MYT1-KD PSIs, likely due to the inclusion of dying cells in these samples. Yet, we could not detect significant MYT1 downregulation in beta cells using scRNA-seq, likely because dying cells were excluded. Future studies of the death-regulating genes in beta cells are needed. Third, how ST18 regulates Ca^2+^ influx is unclear. ST18-KD deregulated the expression of several Ca^2+^, K^+^, and Na^+^ channel-coding genes, likely contributing to the defective glucose-induced Ca^2+^ influx. Yet KCl-induced depolarization did not significantly alter Ca^2+^ influx, suggesting that ion channels (especially Ca^2+^ channels) alone cannot explain the defective Ca^2+^ influx. Other Ca^2+^ regulating processes controlled by ST18 need to be explored in the future. Lastly, the direct transcriptional targets of MYT1 and ST18 are unknown, which need to be addressed in the future using chromatin-binding-based studies.

## Supporting information

ESM description and Figures

ESM Tables

## Author contributions

RH, MY, XT, and GG performed human PSI production, GSIS, IF assays, and imaging. NH did T2D association studies and beta-cell IF imaging. RH performed ITT, and TD performed PSI transplantation. JL and CH helped with viral production and PSI growth. YW and QL analyzed scRNA-seq data, and YX, AJS, and KSL performed scRNA-seq using InDrop. PD and DJ did Ca2+ recording. AB performed human islet isolation. RS, IK, QL, KC, and GG conceptualized the study. All authors participated in manuscript writing/proofing. GG is the guarantor of this work.

## Islet donors

We extend our deepest gratitude to the courageous families who generously donated their loved one’s organs and tissues for biomedical research. Such necessary research as this would not be possible without this selfless gift of hope. Thanks to the efforts of IIDP and Network for Hope (Louisville and Cincinnati) and Lifeline of Ohio, Columbus, for supporting these special families and providing human research pancreases to Dr. Balamurugan’s islet lab. We also thank members of Dr. Balamurugan’s islet lab for their contributions to islet research and transplant.

