## Supplementary material for "Diabetes-associated *MYT1* and *ST18* regulate human beta-cell insulin secretion and survival via other diabetes-risk genes": ESM description and Figures

**Summary of Electronic Supplementary Materials (ESMs):**

1) Materials and Methods.
2) ESM table description (one file with 17 Tabs).
3) ESM Figures and legend (six figures).
4) Human-Islet-Checklist.

**Materials and Methods:**

**Cell culture, transfection, and gene expression assays**

The HEK293 culture followed published protocols. DNA construct transfection using routine methods with Lipofectamine (MIRUS). To compare eGFP levels, images were captured under identical conditions across all samples. The fluorescence intensity was then compared using ImageJ.

**Antibody staining**

Antibody staining was performed using routine methods, with cells spun onto slides or tissue sections. Rabbit anti-Tom20 was from Abcam. Other antibodies were the same as those in the manuscript.

**Mitochondrial and insulin vesicle assays**

Human islet cells attached to glass-bottom plates were infected with lentivirus expressing control and targeting shRNAs. Five days later, cells were fixed and stained with antibodies against insulin and Tom20. Super-resolution images were acquired with Leica-880 LSM or Nikon-NSPARC optics (z-stacks, ~0.2 μm per slice) and quantified using ImageJ. To examine mitochondrial transmembrane potential, PSIs with KD were dissociated into single cells and incubated with MitoView-633 for flow cytometry-based assays. MItoView 633 is a dye that measures mitochondrial transmembrane potential (PMC10627182/PMID: 37936577).

**Real-time RT-PCR assays**

Real-time RT-PCR assays follow those described in the manuscript. Oligos used were: for plasmid DNA copy number: 5'-aactttggcattgtggaagg-3'+ ' + 5'-ggatgcagggatgatgttct-3'; for mCherry: 5'-aactttggcattgtggaagg-3'+ ' + 5'-ggatgcagggatgatgttct-3.

**Supplementary Table. Spreadsheet for gene expression analysis.** Presented as a single Excel file includes 17 tabs.

**ESM Table 1:** Complete list of genes with detectable expression in human PSIs.

**ESM Table 2:** Complete list of genes with detectable expression in human beta cells.

**ESM Table 3:** PSI-expressed T2D-associated genes (GWAS catalog).

**ESM Table 4:** Beta-cell-expressed T2D-associated genes (GWAS catalog).

**ESM Table 5:** PSI-expressed T2D-associated genes (Suzuki list).

**ESM Table 6:** Beta-cell-expressed T2D-associated genes (Suzuki list).

**ESM Table 7:** PSI-expressed T2D-altered genes (Walker list).

**ESM Table 8:** Beta-cell-expressed T2D-altered genes (Walker list).

**ESM Table 9:** DEGs between control and *MYT1*-KD PSIs.

**ESM Table 10:** DEGs between control and *ST18*-KD PSIs.

**ESM Table 11:** DEGs between *ST18*-KD and *MYT1*-KD PSIs.

**ESM Table 12:** Pathways/processes deregulated in pairwise comparisons between *MYT1*-KD, *ST18*-KD, and controls from Deseq2.

**ESM Table 13:** Deregulated genesets in *MYT1*-KD vs control PSIs.

**ESM Table 14:** Deregulated genesets in *ST18*-KD vs control PSIs.

**ESM Table 15:** Deregulated genesets in *ST18*-KD vs *MYT1*-KD PSIs.

**ESM Table 16:** DEGs between control and *MYT1*-KD single beta cells.

**ESM Table 17:** DEGs between control and *ST18*-KD single beta cells.

**Supplementary Figures:**

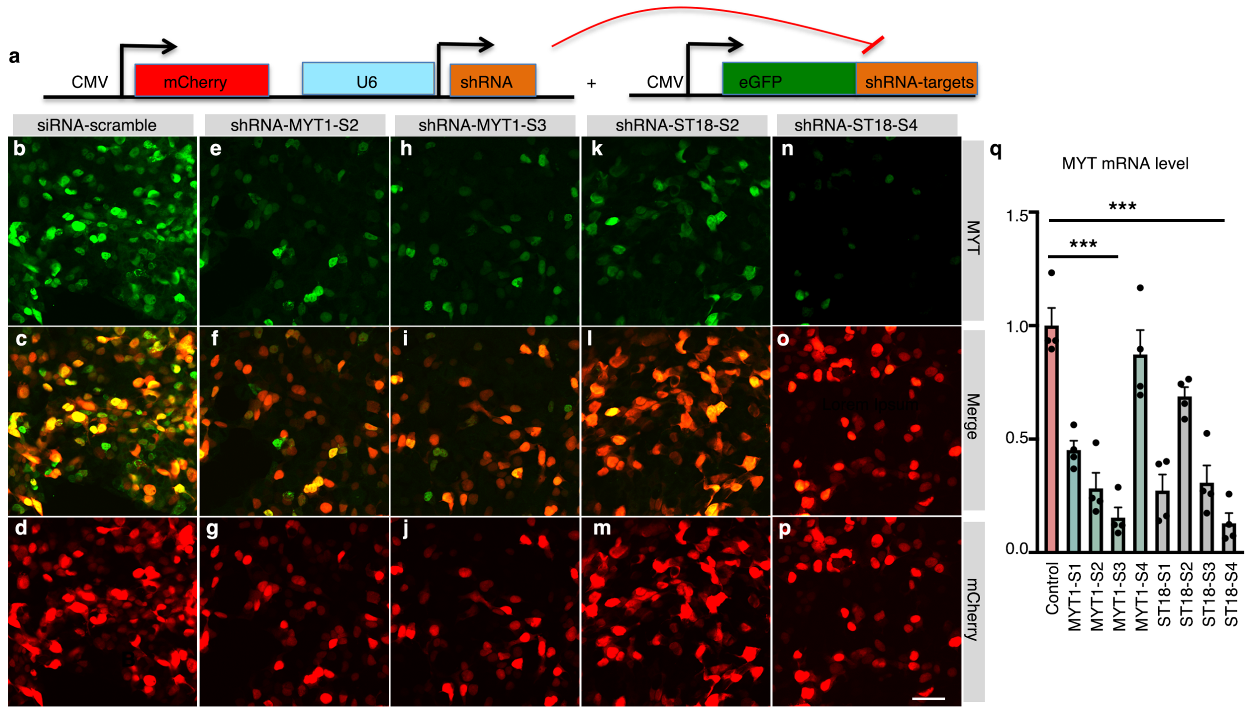


**ESM Fig. 1. Identification of shRNAs for human *MYT1* and *ST18* KD.** Four shRNA sequences were tested for each gene. (a) A diagram showing the constructs to test the efficacy of shRNA. (b-p) Reporter assays showing the different efficacy of shRNA KD of reporter eGFP. Note that images of two MYT1 shRNA results (MYT1-S2 and MYT1-S3) and two ST18 shRNA results (ST18-2 and ST18-4) were included. (q) RT-PCR results to examine the mRNA downregulation by all eight targeting shRNAs. The control was a CMV-driven scrambled RNA. Bar, 20 μm.

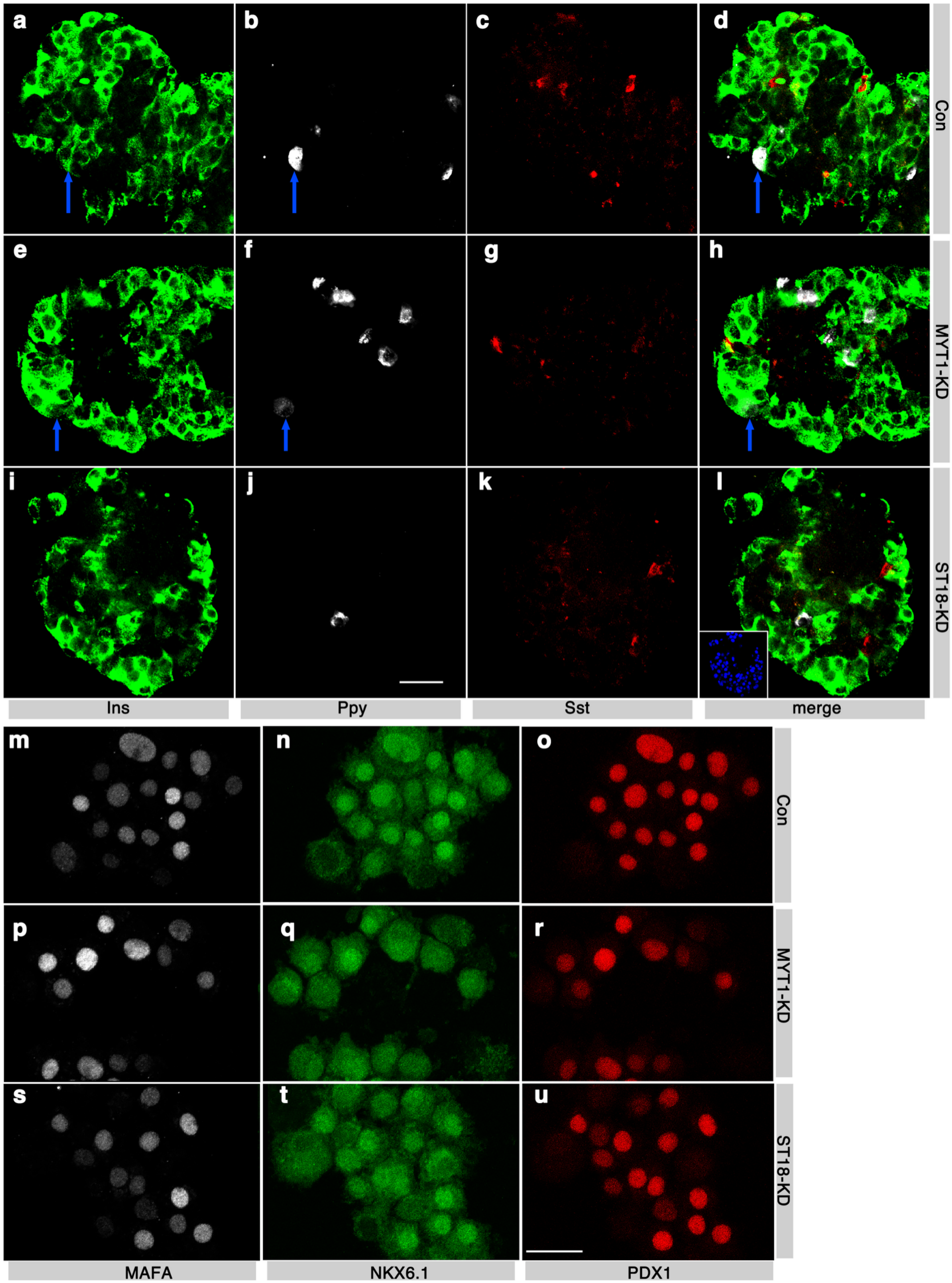


**ESM Fig. 2. *MYT1*- or *ST18*-KD does not induce co-expression of islet hormones or alter the expression of several TFs in β cells.** (a-l) Insulin co-expression with pancreatic polypeptide (PPY) and somatostatin (SST) in freshly prepared PSIs. Inset in l is a DAPI staining to locate all nuclei. (m-u) MAFA, NKX6.1, and PDX1 expression in control and KD-ed cells. (a-i) Single optical slices were shown. (m-u) Z-projections of the entire cell were shown. Blue arrows in a-d or e-h point at two cells co-expressing INS and PPY. Bars, 20 μm.

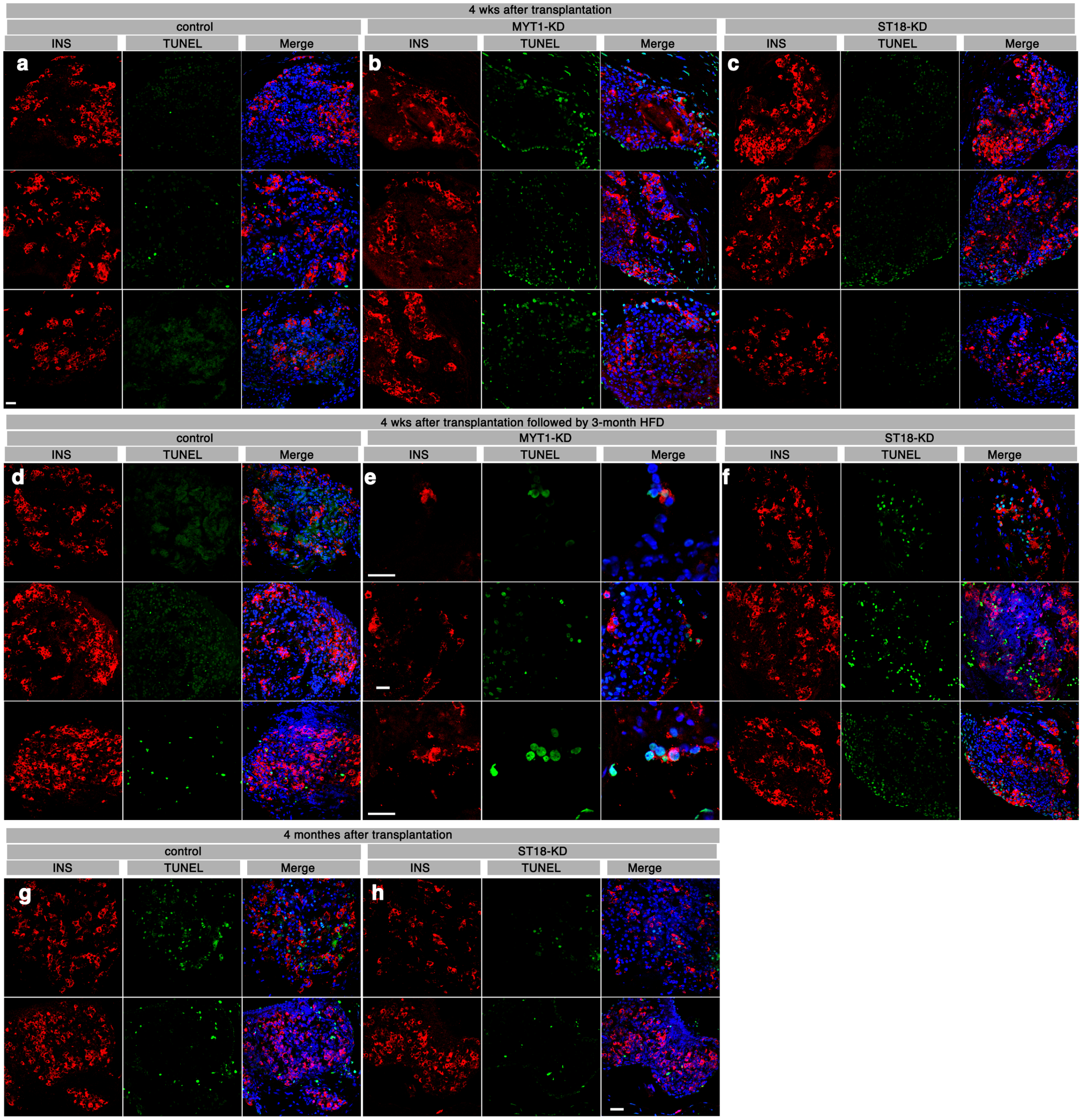


**ESM Fig. 3. Extra images of β-cell death with *MYT1*- or *ST18*-KD.** (a-c) TUNEL staining in transplanted PSIs 4 weeks after transplantation. Each row represented PSIs from one donor. Single channels of insulin, TUNEL, and merged images (with DAPI) were included. (d-f) Same as a-c, except **that** PSIs were used after 12 additional weeks of HFD challenge. (g and h) Same as d-f except HFD was not used. Note that in g and h, nearly all green cells are not $INS^{+}$ (please zoom in to inspect).

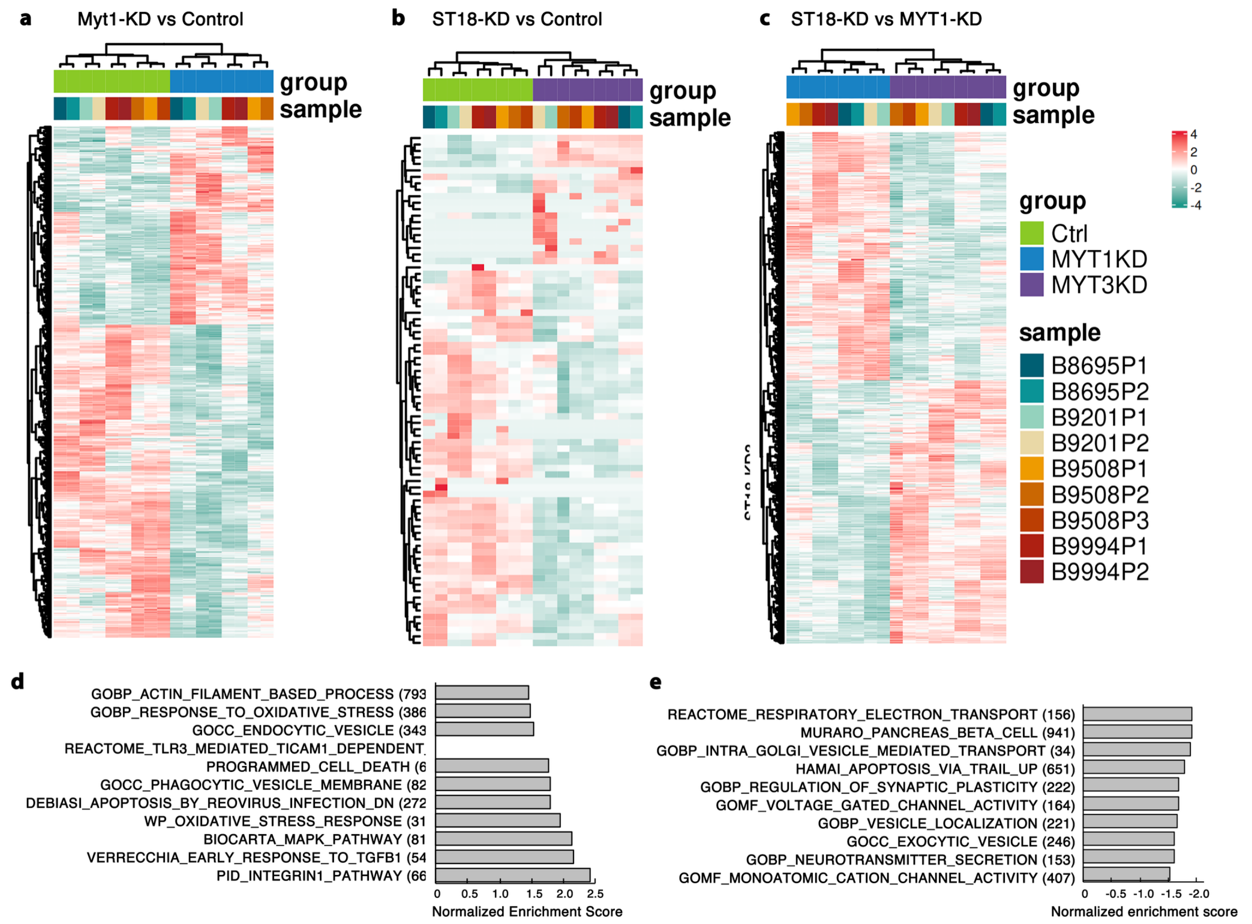


**ESM Fig. 4. DEGs and changed genesets in pairwise comparison between control, *MYT1*-KD, and *ST18*-KD PSIs.** (a-c) Hierarchical clustering of DEGs (from Deseq2) between control–*MYT1*-KD (a), control–*ST18*-KD (b), and *MYT1*-KD–*ST18*-KD samples (c). Technical replicates for each donor were included (the same designation corresponds to those deposited RNA-seq raw data). (d and e) Up- or down-regulated genesets in *ST18*-KD over *MYT1*-KD PSI samples. Ten examples of each category were shown (see ESM Table 15 for the complete list).

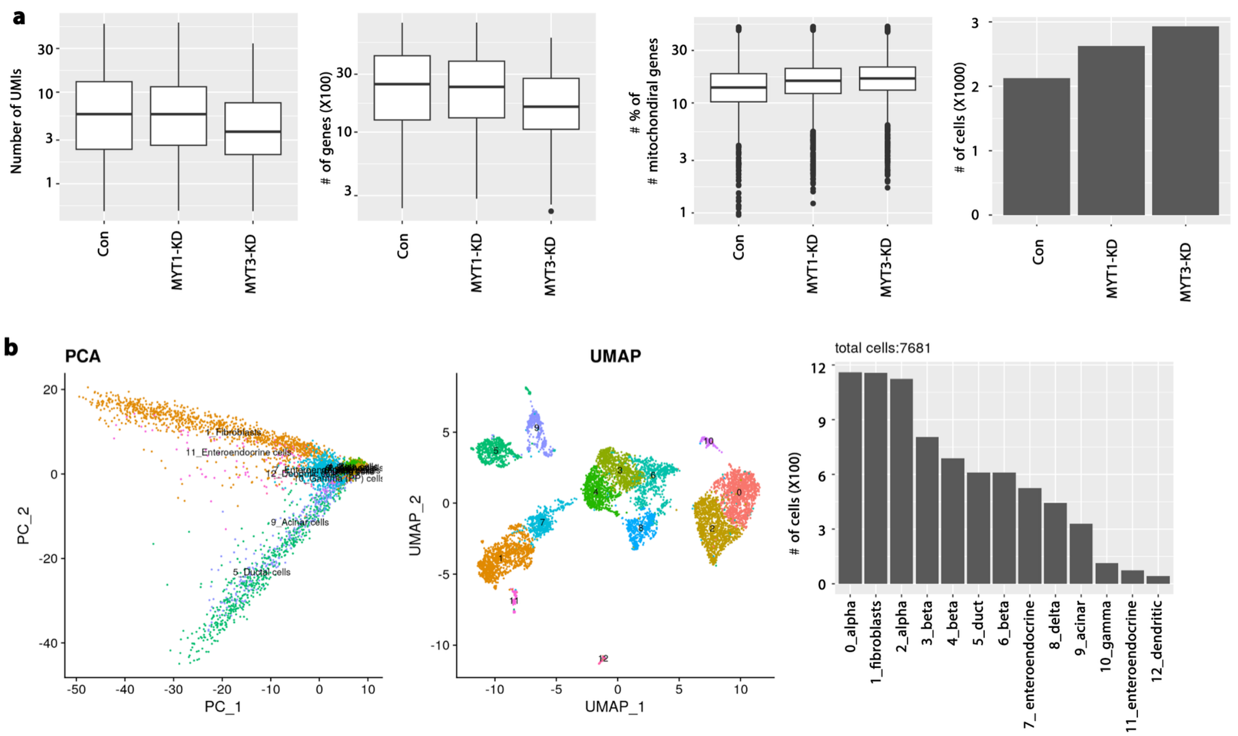


**ESM Fig. 5. Quality controls showing the sc-RNAseq for one batch of human PSI (control, *MYT1*-KD, or *ST18*-KD).** (a) The number of UMIs, genes (total or mitochondrial specific), or cells in each sample. (b) Principal-component plots, or UMAPs, and bar graphs showing the distribution and numbers of different islet cell types.

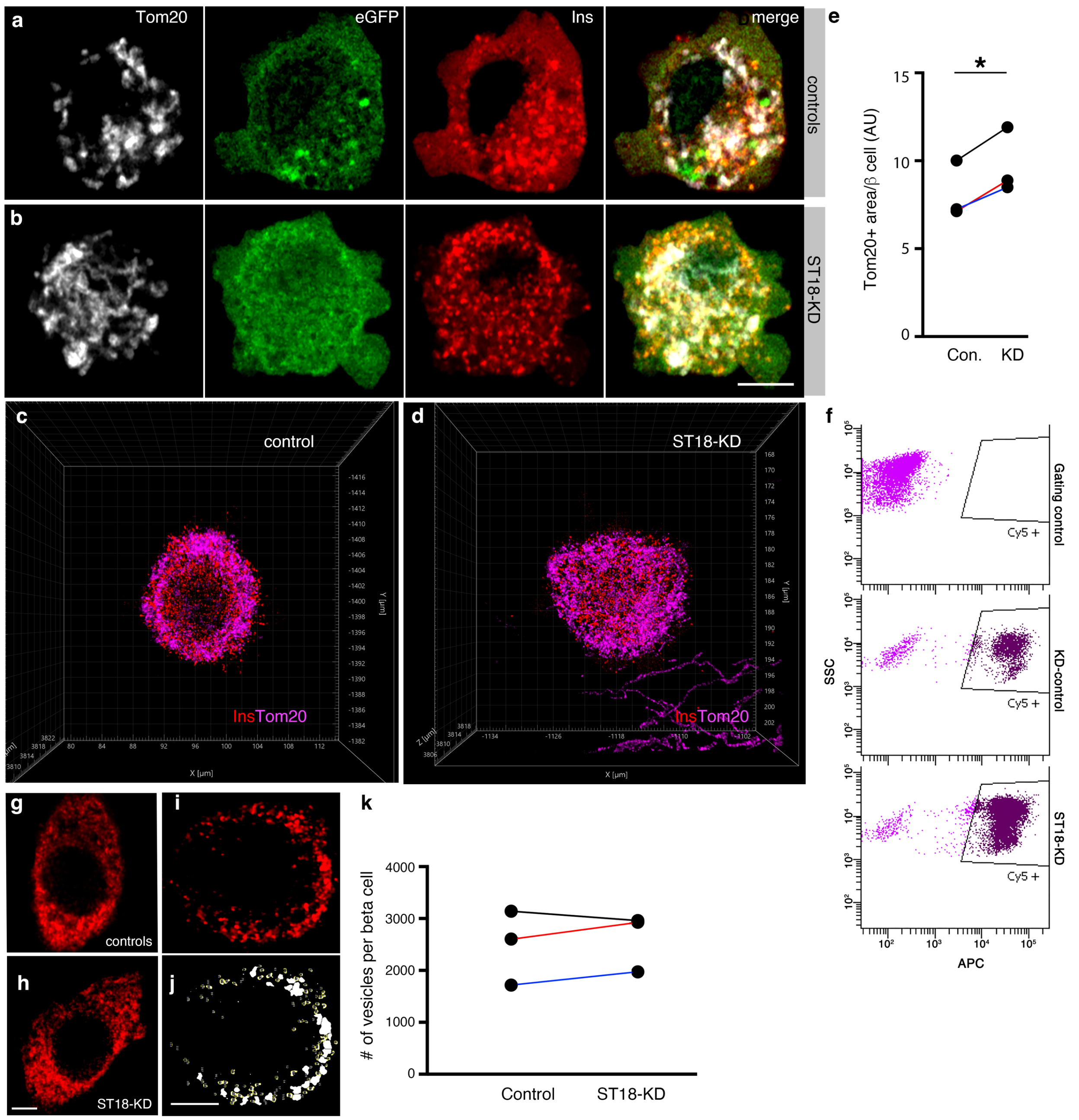


**ESM Fig. 6. *ST18*-KD increased mitochondrial volume but did not affect mitochondrial transmembrane potential and insulin granule production.** (a-e) Mitochondrial volume in each β cell after *ST18*-KD. (a and b) Single slice of confocal images (white: Tom20, mitochondrial membrane; green, shRNA expression; red, insulin). (c and d) 3-D projections (IMARIS) of insulin granules and mitochondria in control and *ST18*-KD beta cells. (e) Quantification of mitochondrial volume in each beta cell, represented by the total mitochondrial area on all optimal slices. Results from three donors were presented, with those from the same donor connected by lines of different colors (black: donor #5, red: donor #6; blue: donor #7, see Human-Islet-Checklist). Ten to 14 beta cells were counted for each sample (donor/genotype). *P* is from repeated measure ANOVA. *: $p<0.05$. (f) A typical flow cytometry assay of transmembrane potential in mitochondria of PSI cells (indicated by the Cy5 intensity). Note the unequal distribution of gated cells, likely reflecting the different islet cell types in PSI. (g-k) Quantification of the number of vesicles in each beta cell. (g and h) Typical insulin vesicles in control and *ST18*-KD beta cells. (I and j) Quantification method, with the non-contacting vesicles counted by ImageJ and contacting vesicles counted manually (j). Beta cells from three donors were counted. For each donor, 10-17 beta cells were examined. All optical slices that were 0.88 micrometers apart were counted to avoid counting vesicles multiple times. In k, the combined results of three donors were presented, with those of the same donor connected by lines of different colors (black: donor #7, red: donor #5, blue: donor #6). Bar, 5 μm.

**Human-Islet-Checklist**

### Checklist for reporting human islet preparations used in research

Adapted from Hart NJ, Powers AC (2018) Progress, challenges, and suggestions for using human islets to understand islet biology and human diabetes. Diabetologia https://doi.org/10.1007/s00125-018-4772-2

| **Islet preparation** | **1** | **2** | **3** | **4** | **5** | **6** | **7** |
|---|---|---|---|---|---|---|---|
| Unique identifier | **RP-003** | **SAMN31645455** | **SAMN37350251** | **SAMN28157682** | **SAMN34130383** | SAMN49948188 | SAMN50225209 |
| Donor age (years) | 48 | 68 | 43 | 37 | 42 | 54 | 53 |
| Donor sex (M/F) | F | M | F | M | F | M | F |
| Donor BMI (kg/m$^2$) | 22.3 | 30.7 | 29.9 | 30.3 | 29.3 | 31.6 | 33.4 |
| Donor $HbA_{1c}$ or other measure of blood glucose control | 5.0 | 5.4 | 5.2 | 5.0 | 5.2 | 5.8 | 5.8 |
| Origin/source of islets[b] | NORTON ISLET CENTER | IIDP | IIDP | IIDP | IIDP | IIDP | IIDP |
| Islet isolation centre | NORTON ISLET CENTER | Prado | Wisconsin | SC-ICRC | Prado | Imagine islet Center | SC-ICRC |
| Donor history of diabetes? Please select yes/no from drop down list | No | No | No | No | No | No | No |
| Diabetes duration (years) | NA | NA | NA | NA | NA | NA | NA |
| Glucose-lowering therapy at time of death[c] | NA | NA | NA | NA | NA | NA | NA |
| Donor cause of death | Head Trauma | Anoxia | Head trauma | stroke | *Cerebrovascular/stroke* | *Cerebrovascular/stroke* | *Cerebrovascular/stroke* |
| Warm ischaemia time (h) | 9 minutes | None | None | None | None | None | 21 minutes |
| Cold ischaemia time (h) | 9 hours 8 mins | 10 hours 20 minutes | 5 hours | 7 hours 20 minutes | none | 2 hour 50 minutes | 9 hour 53 minutes |
| Estimated purity (%) | 90 | 85 | 90 | 80 | 90 | 85 | 95 |
| Estimated viability (%) | 95 | 90 | 95 | 96 | 90 | 90 | 95 |
| Total culture time (h)[d] | 60 hours | 72 hours | 92 hours | 96 hours | 118 hours | 64 hours | 46 hours |
| Glucose-stimulated insulin | GSIS, Stimulation | GSIS, SI is 3.5 | GSIS, SI 3.4 | GSIS, SI is 2.3 | GSIS, SI, 3.2 | GSIS, SI: 2.6 | GSIS, SI: 3.1 |

| | | | | | | | |
|---|---|---|---|---|---|---|---|
| secretion or other functional measurement[e] | index (SI) is 2.1 | | | | | | |
| Handpicked to purity? Please select yes/no from drop down list | Yes | Yes | Yes | Yes | Yes | Yes | Yes |
| Additional notes | For Ca2+ recording, mitochondrial studies, and ISG studies. | Used in Bulk-RNAseq and transplantation | Bulk-RNAseq, scRNAseq and transplantation | Used in Bulk RNAseq | Used in Bulk-RNAseq, transplantation | For Ca2+ recording, mitochondrial studies, and ISG studies. | For Ca2+ recording, mitochondrial studies, and ISG studies. |

[a]If you have used more than eight islet preparations, please complete additional forms as necessary
[b]For example, IIDP, ECIT, Alberta IsletCore
[c]Please specify the therapy/therapies
[d]Time of islet culture at the isolation centre, during shipment and at the receiving laboratory
[e]Please specify the test and the results
